# *In vivo* prenylomic profiling in the brain of a transgenic mouse model of Alzheimer’s disease reveals increased prenylation of a key set of proteins

**DOI:** 10.1101/2022.04.01.486487

**Authors:** Angela Jeong, Shelby A. Auger, Sanjay Maity, Ling Li, Mark D. Distefano

## Abstract

Dysregulation of protein prenylation has been implicated in many diseases, including Alzheimer’s disease (AD). Prenylomic analysis, the combination of metabolic incorporation of an isoprenoid analogue (C15AlkOPP) into prenylated proteins with a bottom-up proteomic analysis, has allowed identification of prenylated proteins in various cellular models. Here, transgenic AD mice were administered with C15AlkOPP through intracerebroventricular (ICV) infusion over 13 days. Using prenylomic analysis, 36 prenylated proteins were enriched in the brains of AD mice. Importantly, the prenylated forms of 15 proteins were consistently upregulated in AD mice compared to non-transgenic wild-type controls. These results highlight the power of this in vivo metabolic labeling approach to identify multiple post-translationally modified proteins that may serve as potential therapeutic targets for a disease that has proved refractory to treatment thus far. Moreover, this method should be applicable to many other types of protein modifications, significantly broadening its scope.

## Main

Advances in science and medicine over the last 100 years have triggered a dramatic increase in life expectancy. With a longer-living population, it is expected that there will be an increase in aging-related diseases including Alzheimer’s disease (AD). This debilitating diagnosis is not exclusive in its devastation, and it places the family of the afflicted under extreme emotional and financial stress. Currently there are over 6 million Americans living with AD, and by 2060 that number is projected to grow to 13.8 million.^1^ The traditional pathological markers of AD, including aggregation of amyloid-β peptide (Aβ) in neuritic plaques and formation of neurofibrillary tangles from hyper-phosphorylation of Tau protein, have been meticulously characterized and pursued as potential targets for therapeutic development.^2–4^ Unfortunately, which dysregulated biological mechanisms lead to the pathogenesis of AD, especially sporadic AD, is still not well understood. In the search for possible mechanisms for AD pathogenesis, one important process, that has been implicated yet underexplored, is protein prenylation.

Protein prenylation is a widespread post-translational modification of proteins consisting of the addition of an isoprenoid near the C-terminus for intracellular protein localization and trafficking.^5–8^ The prenylation of proteins is integral to proper cellular signaling and regulation and specific prenylated proteins, including Rab10 and H-Ras, have been associated with the development of AD.^6,9–12^ There are three types of prenylation: farnesylation, type I geranylgeranylation, and type II geranylgeranylation. During farnesylation and type I geranylgeranylation, either a farnesyl group from farnesyl diphosphate (FPP, Fig.1A) or a geranylgeranyl group from geranylgeranyl diphosphate (GGPP, Fig. 1A) molecule is transferred to a protein via the action of farnesyltransferase (FTase) or geranylgeranyltransferase type I (GGTase I) respectively.^13^ Protein substrates for farnesylation or type I geranylgeranylation have an identifying tetrapeptide motif at the C-terminus recognized by these enzymes. This CaaX motif, where C is cysteine and the site of modification, a is an aliphatic amino acid, and X is a variable amino acid, determines whether a protein is prenylated and which type of prenylation occurs.^14^ For type II geranylgeranylation, there are different motifs, CXC, XCXC, and CC, which have two cysteine residues for the transfer of two geranylgeranyl groups from GGPP. An additional upstream sequence element is required for this type of modification. The substrate scope for geranylgeranylation type II is exclusive to Rab proteins, a subset of the small GTPase proteins.^15^

**Figure 1.**
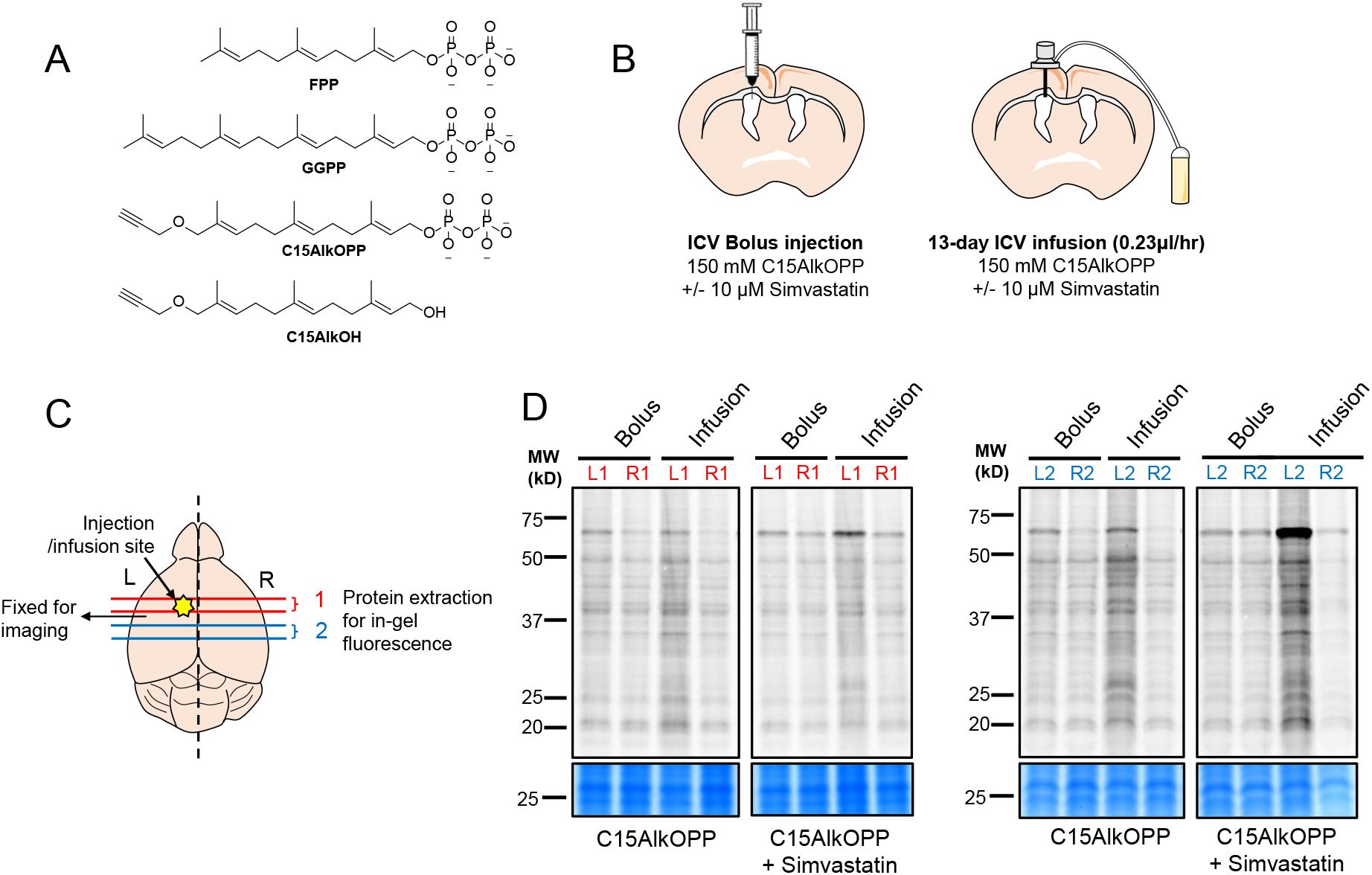
Brain metabolic labeling after a single bolus ICV injection and 13-day ICV infusion of C15AlkOPP. (**A**) Structures of endogenous substrates for prenylation, alkyne-containing diphosphate analogue (C15AlkOPP) used in this study and the alcohol (C15AlkOH) analogue. (**B**) Schematic representation of ICV bolus injection and ICV infusion and (**C**) the relative locations of brain coronal sections (400 μm) from the injection/infusion site that were used for the representative in-gel fluorescence. The brain section (600 μm) located between section #1 and #2 was PFA-fixed for *in situ* click reaction and imaging. (**D**) In-gel fluorescence and Coomassie blue gel staining images of brain regions depicted in B. Brains were harvested 48 hours after the left ICV bolus injection of 150 mM C15AlkOPP (10 μL) or 10 mM Simvastatin (SV) (10 μL) + 150 mM C15AlkOPP (10 μL); or 13 days after the initiation of left ICV infusion of followings: 150 mM C15AlkOPP, or 10 mM simvastatin (SV) + 150 mM C15AlkOPP (1:3).

Prenylation was initially linked to AD through retroactive epidemiological analysis of clinical records that demonstrated that patients who take statins have significantly lower incidences of AD diagnosis.^4,10,16–18^ Statins, which are 3-hydroxy-3-methylglutaryl coenzyme A (HMG-CoA) reductase inhibitors, decrease cholesterol production by suppressing the production of mevalonate and the biosynthesis of the downstream isoprenoids FPP and GGPP. This decrease in isoprenoid production, and concomitant decrease in protein prenylation, is likely the mechanism behind statin-induced, cholesterol-independent pleiotropic effects including neuroprotection.^4,17,19^ Beyond epidemiological data, there is also compelling biochemical evidence that directly shows that prenylation is dysregulated in AD brains. Analysis of brain tissue from AD patients has revealed increased levels of FPP and GGPP and elevated mRNA expression for the corresponding synthases for these two compounds.^20^ Recent studies further showed that FTase per se, its farnesylated substrates, and downstream signaling pathways were significantly increased in human AD brains.^12^ This dysregulation, was further confirmed in experiments using the transgenic AD model APP/PS1 mice, in which either FTase haplodeficiency or neuron-specific FTase-deficiency rescued cognitive function, decreased Aβ deposition, and attenuated neuroinflammation.^12,21^ Interestingly, GGTase haploid deficient APP/PS1 mice showed a decrease in Aβ and neuroinflammation but without rescuing cognitive function, highlighting potential distinct roles of the different types of prenylation in AD.^22^ While some prenylated proteins have been individually implicated in the development of AD and Aβ processing including H-Ras, Rac1, Rho, Rab5, Rab7, Rab10, and Rab35,^9,12,23–27^ a method for simultaneously tracking the prenylation of all prenyltransferase protein substrates would be highly useful for clarifying the role of protein prenylation in AD.

Prenylomic analysis involves the use of metabolic labeling of prenylated proteins with an alkyne-containing analogue of FPP and GGPP (Fig. 1A), bioorthogonal labeling and subsequent enrichment followed by quantitative bottom-up proteomic analysis. Metabolic labeling is possible since FTase, GGTase I, and GGTase II have shown some flexibility with regards to the isoprenoid substrate structure, allowing for the development of analogues of the natural substrate containing biorthogonal functionality. The structure of C15AlkOPP, an analogue that is an efficient substrate for both FPP- and GGPP-utilizing prenyltransferases is shown in Fig. 1A. ^28–30^ Prenylomic analysis using C15AlkOPP, and related compounds, have been employed to identify and track the levels of specific prenylated proteins in cell culture-based systems and to explore their roles in diseases including cancer.^31–34^ In earlier work, this method was used to profile prenylated proteins in brain-derived cell lines using both primary and immortalized cells.^35^ Those experiments revealed similarities and differences in the identities and levels of prenylated proteins found between different cell types. However, such experiments using only one cell type cannot capture the complex pathology of a disease such as AD. While metabolic labeling in mice has been previously reported,^36^ the application of metabolic labeling combined with proteomic analysis is quite limited.^37^

Here, methodology is described that extends the application of prenylomic analysis to the transgenic APP/PS1 mouse model that recapitulates many of the hallmarks of AD. First, different methods for delivery of C15AlkOPP to the brain of non-transgenic wild-type (WT) mice were explored to maximize incorporation of the analogue. It was found that constant slow intracerebroventricular (ICV) infusion of C15AlkOPP for approximately two weeks using an osmotic pump gave substantially higher levels of protein labeling compared with single injections as determined via in-gel fluorescence analysis. Using this methodology, three pairs of APP/PS1 and WT mice were labeled with C15AlkOPP, followed by prenylomic analysis. The triplicate comparison of APP/PS1 to WT mice showed a consistent increase in prenylation in the AD model. A group of 15 prenylated proteins were identified across all three pairs. Importantly, many of these proteins have a documented relationship to the neuropathology of AD. This is the first example of the use of prenylomic analysis to characterize the dysregulation of a prenylome *in vivo*.

## Results

### A single bolus ICV injection of C15AlkOPP showed limited brain labeling of prenylated proteins

Previously, brain metabolic labeling was attempted by intraperitoneally (IP) injecting an isoprenoid analogue C15AlkOH (Fig. 1A) dissolved in normal saline containing 5% Tween 80. Although IP delivery of C15AlkOH successfully labeled prenylation substrates in the peripheral organs, it failed at brain labeling, possibly due to its limited blood brain barrier (BBB) permeability (unpublished data). Therefore, a direct brain injection approach was employed to ensure the delivery of the isoprenoid analogue to the brain. Diphosphate isoprenoid analogue C15AlkOPP manifests superior labeling efficiency compared to alcohol based C15AlkOH in multiple cell lines^38^ and therefore, C15AlkOPP was chosen over C15AlkOH in this study. The chemical structures of C15AlkOPP, C15AlkOH, and the endogenous isoprenoids FPP and GGPP are shown in Fig. 1A.

To maximize the potential injection volume and brain distribution, the lateral ventricle was chosen as the target site of administration (Extended Data Fig. 1A). After unsuccessful attempts using 10 mM C15AlkOPP, the injection concentration of C15AlkOPP was increased to 150 mM. After 48 hours following a 10 µL bolus left ICV injection of vehicle or 150 mM C15AlkOPP with or without co-administration of 10 mM simvastatin (SV), brains were collected from mice. To survey the distribution of the analogue in relation to the injection site, both left and right cerebral hemispheres were sliced at alternating thickness of 400 μm and 600 μm. Brain slices of 600 μm thickness were fixed for in situ click reaction. Meanwhile, all the 400-μm slices that were located within 2 mm anterior or posterior to the injection site were homogenized and subjected to click reaction with TAMRA-azide for visualization of metabolic labeling. In-gel fluorescence showed increased TAMRA fluorescence in the 60-65 kDa region in mice injected with C15AlkOPP compared to mice injected with vehicle, in the brain region adjacent to the injection site (Extended Data Fig. 1B and 1C). This labeling was greater in the mouse that received the mixture of both SV and the analogue because statins increase the incorporation efficiency of exogenous analogues into target proteins by reducing endogenous FPP and GGPP levels.^28,35,39^ However, surprisingly, there was no TAMRA fluorescence band in the 20-25 kDa region of the gel where many prenylated small-GTPases migrate, suggesting that C15AlkOPP did not efficiently label small GTPases in this experiment (Extended Data Fig. 1 C).

To verify that the specific analogue solution used for the injection could modify small GTPases and other known prenylation substrates, the same C15AlkOPP solution was diluted and tested using the human neuroblastoma SH-SY5Y cell line at a final concentration of 10 μM. After 24 hours, the incorporation of the analogue was visualized using in-gel fluorescence. The protein labeling pattern in SH-SY5Y cells was similar to that previously reported in HeLa and COS-7 cells^28,35^ with intense fluorescence bands in the 20-25 kDa region (Extended Data Fig. 1D). These results confirmed the integrity of the analogue sample and that the observed absence of small GTPases labeling *in vivo* most likely resulted from poor labeling efficiency due to the single bolus injection approach.

### *In vivo* brain labeling with sustained ICV infusion of C15AlkOPP

To achieve prolonged delivery of the isoprenoid analogue, osmotic pump mediated ICV infusion was employed. After 13 day-infusions of 150 mM C15AlkOPP, or a 1:3 mixture of 10 µM SV and 150 mM C15AlkOPP to the left ventricle (Fig. 1A, right diagram), brains were collected and sectioned the same way as described for the bolus ICV injection samples. Fluorescent labeling of lysate obtained from each 400 μm slice proximal to the infusion showed intense fluorescent signals throughout the entire lane including the 20-25 kDa region (Fig. 1B). As expected, co-administration of SV with C15AlkOPP led to more intense labeling than infusing C15AlkOPP alone, but the presence of SV did not change the overall fluorescent banding pattern. In addition, the labeling was restricted to the brain regions ipsilateral to the injection/infusion site (left). To directly visualize the distribution and cellular uptake of C15AlkOPP following the 13-day infusion in the brain, prefixed 600 μm brain slices from the region adjacent to the most intensely labeled brain slides per in-gel fluorescence were selected. These slides were further sectioned at a thickness of 50 μm, and in situ click reaction was performed directly on the section using TAMRA-azide along with the immunostaining of a neuronal marker NeuN (Extended Data Fig. 2). Even though significant background labeling was detected, overall, the TAMRA fluorescence intensity in the C15AlkOPP-treated brain section was significantly higher than that in the vehicle-treated mouse brain section. This supports the conclusion that C15AlkOPP was successfully delivered to the brain parenchyma around the infusion site after being introduced into the ventricle.

### Prenylomic profiling of brains from WT mice following ICV infusion of C15AlkOPP with or without statin

To identify and quantify the prenylated proteins that were visualized in both the in-gel fluorescence analysis (Fig. 1D) and in-situ labeling experiments (Extended Data Fig. 2), the prenylomic workflow shown in Fig. 2A was employed. This allows the identities and relative levels of prenylated proteins to be determined through a combination of metabolic incorporation of C15AlkOPP, bioorthogonal labeling, and subsequent enrichment followed by quantitative bottom-up proteomic analysis. Initial profiling experiments were performed comparing a WT mouse that had undergone a 13-day ICV infusion with a mouse that received the probe vehicle in the same manner. For the enrichment of these samples, the brain lysate containing 2 mg of total protein from each mouse was subjected to CuAAC labeling with biotin-azide.

**Figure 2:**
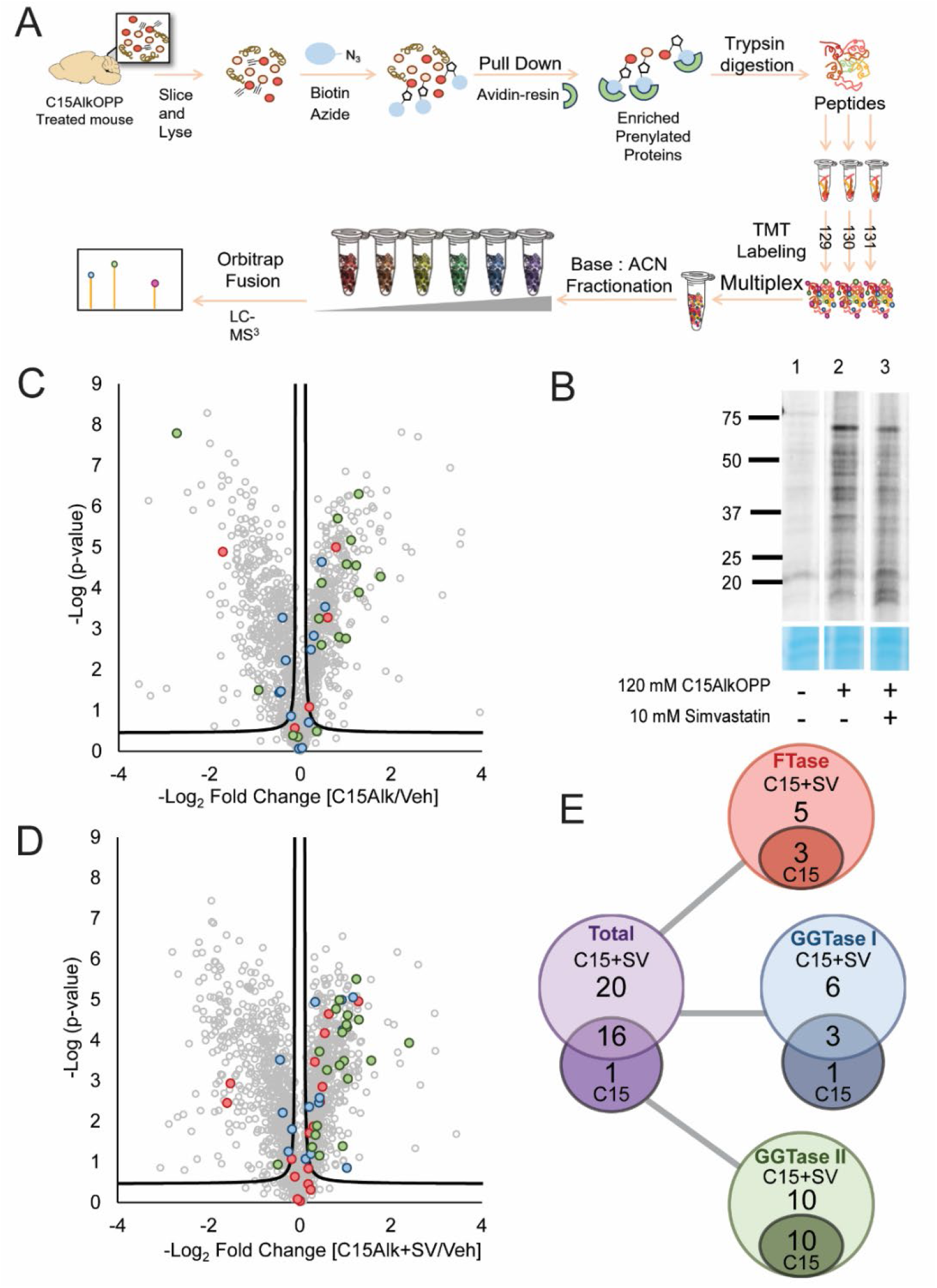
Prenylomic profiling of WT mice after ICV infusion of C15AlkOPP with or without a statin treatment. (**A**) Schematic representation of the prenylomic workflow depicting the enrichment of prenylated proteins, their digestion, and the subsequent division of peptides into three replicates. This scheme shows the process for C15AlkOPP treated samples. The vehicle sample was treated in the same manner with the only change being that vehicle replicates were labeled with TMT 126, TMT127, and TMT128 reagents. (**B**) Gel fluorescence analysis of the brain samples used in this analysis. After harvesting, the left side of the brain was lysed and subjected to a click reaction with TAMRA-azide to visualize the level of alkyne-probe incorporation in these samples. Lane 1 contains vehicle sample, Lane 2 contains C15AlkOPP sample and Lane 3 contains the sample treated with C15AlkOPP and statin. (**C**) Volcano plot comparing C15AlkOPP with vehicle ICV infusion (FDR=5%). (**D**) Volcano plot comparing C15AlkOPP and simvastatin coadministration with vehicle ICV infusion (FDR=5%). (**E**) Venn diagrams showing the distribution of prenylated proteins obtained from the volcano plots shown in panels **C** and **D**. These are further subdivided by type of prenylation shown in color. Proteins that are grouped differently across the pairs are entered as one protein. Example: In C15AlkOPP treatment alone Rab3a,3b,3c are grouped, and in the SV+C15AlkOPP treated mouse Rab 3a,3b, and 3c are not grouped. Therefore, for simplicity, Rab 3a,3b and 3c are one data point in the Venn diagram. Color scheme: prenylated proteins that are known substrates for FTase (red); prenylated proteins that are substrates for GGTase I (blue); prenylated proteins that are known substrates for GGTase II (green).

As previously established,^35^ this method allows for the enrichment of prenylated proteins from complex mixtures using a neutravidin-resin pull down followed by on-bead digestion with trypsin. The resulting peptide samples were then divided into three technical replicates, as shown in Fig. 2A, for a TMT 6plex labeling quantification strategy. Before mass spectrometric analysis, the samples were fractionated at high pH and then subjected to a Nano-LC-MS^3^ separation and analysis. The resulting data was processed with MaxQuant using a non-redundant mouse protein reference library.

In this initial proof-of-concept work, the level of the incorporation of C15AlkOPP in the brain of WT mice after the 13-day ICV infusion allowed the detection of 17 prenylated proteins in mice treated with the analogue versus the vehicle (Fig. 2C, SI Table 3). Those 17 proteins are mostly substrates for GGTase I (4 proteins, blue) or II (11 proteins, green). In previous work, performed in cell culture, it has been demonstrated that treatment with a statin leads to increased incorporation of C15AlkOPP, as evidenced by a larger number of proteins identified with greater fold changes as well as by an increase in the identification of farnesylated proteins.^35,38,39^ When WT mice were treated with C15AlkOPP in conjunction with simvastatin, a similar increase was seen with 37 prenylated proteins found, compared with 17 observed in the absence of statin. This 20-protein increase included a significant increase in the number of FTase substrates. Thus, in the presence of simvastatin, 8 FTase substrates, 9 GGTase I substrates and 20 GGTase II substrates were detected (SI Table 3). From these two experiments, two important baselines for the identification of prenylated proteins from the *in vivo* metabolic labeling of mouse brains were defined. In the case of C15AlkOPP treatment alone, the identification of 17 proteins suggested that it should be possible to monitor the levels of those proteins in WT versus diseased mice. In contrast, the 37 proteins detected using C15AlkOPP in the presence of a statin likely defines an upper bound for the number of prenylated proteins that might be studied via this metabolic labeling strategy. Thus, these experiments set the stage for the application of this methodology for the study of protein prenylation in the APP/PS1 mouse model.

### *In vivo* brain labeling of APP/PS1 and WT mice via ICV infusion of C15AlkOPP

Next, to test whether the C15AlkOPP-mediated brain labeling can probe prenylomic changes associated with the amyloid pathology, metabolic labeling was performed in a group of aged female (16-20 months) APP/PS1 mice and their WT controls (n=3 mice/genotype). At this age, APP/PS1 mice have significant amyloid plaque pathology as well as cognitive behavioral deficits.^40^ During the preliminary testing of ICV infusion, some precipitation of C15AlkOPP was noted at the concentration of 150 mM. Therefore, the analogue was diluted to 100 mM to avoid any potential precipitation of C15AlkOPP inside the pump or in the brain. Both groups were administered the same amount of C15AlkOPP through a brain infusion osmotic pump (Fig. 3A). Brains were collected 13 days after initiating the ICV infusion. Based on the labeling pattern observed from the brain slices, the left-brain region around the infusion site (including the area 1-2 mm anterior and posterior to the infusion site) was dissected out (Fig. 3B), homogenized and subjected to click reaction with TAMRA-azide and subsequent in-gel fluorescence (Fig.3C). Interestingly, in all 3 cases, the level of fluorescent labeling in the samples obtained from the APP/PS1 mice was substantially higher than what was observed in the samples from the WT mice. This finding suggests that protein prenylation is increased in the brain of AD mice.

**Figure 3.**
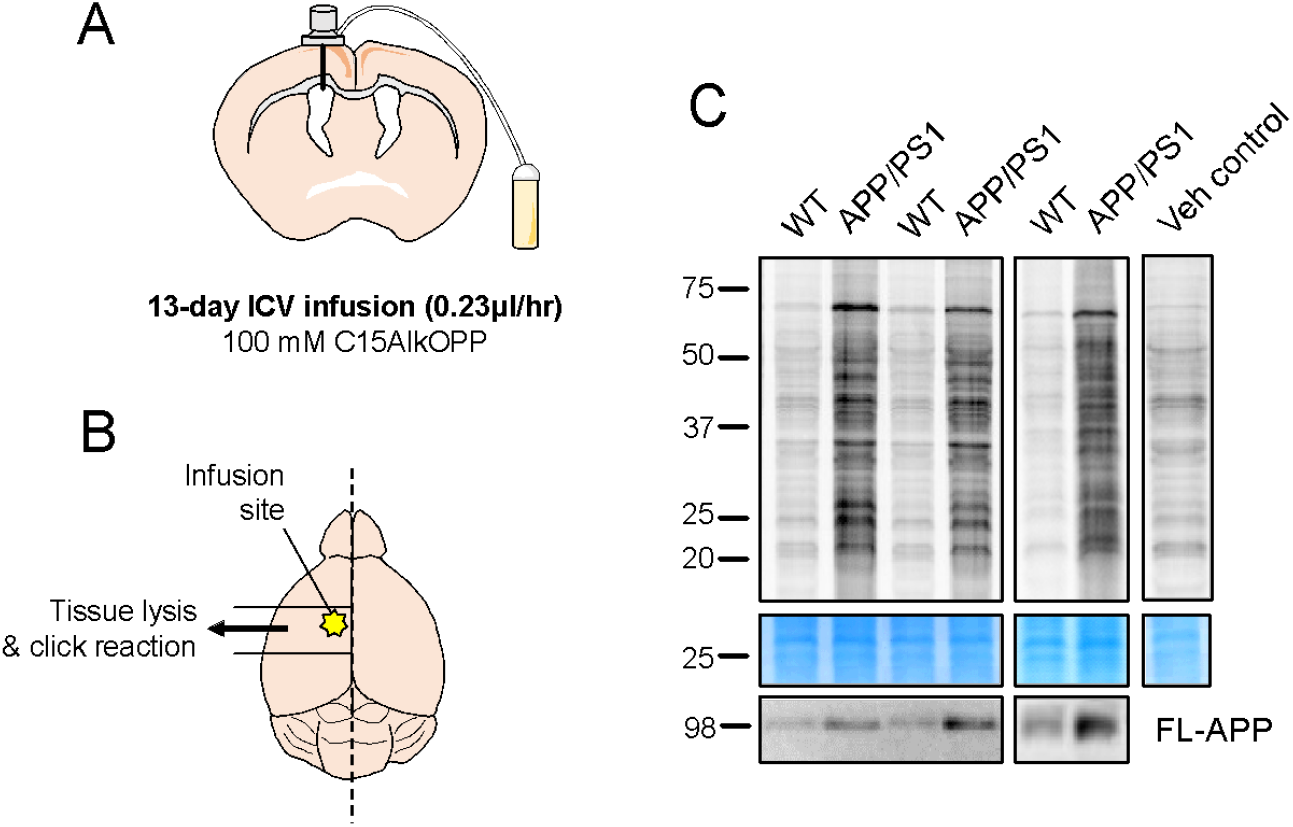
Brain metabolic labeling of APP/PS1 and wild type control mice. (**A**) Schematic representation of the site of ICV infusion and (**B**) brain area used for the representative in-gel fluorescence and the prenylome profiling. (**C**) In-gel fluorescence and Coomassie blue gel staining images of WT and APP/PS1 mice after 13-day left ICV infusion of 100 mM C15AlkOPP. Full-length APP (FL-APP) was measured via immunoblot assay to confirm the genotypes.

### Proteomic analysis identified significantly enriched prenylated proteins in the brain of APP/PS1 mice compared to WT controls

To explore how the levels of prenylation differ in the AD model, the brain samples from three pairs of mice described above were subjected to the prenylomic workflow as illustrated in Fig. 2A. By comparing APP/PS1 mice treated with C15AlkOPP with WT mice treated with the same amount of C15AlkOPP, differences in the levels of specific prenylated proteins in the two groups could be determined. If prenylation is unaffected in AD, then the modified prenylated proteins should not manifest statistically significant fold changes, resulting in them residing in the interior of the volcano plots. In all three pairs this was not the case; instead, it was found across all three pairs that increased prenylation was observed in the AD mice. In total, 26-28 prenylated proteins were enriched preferentially in each of the APP/PS1 mouse vs WT mouse comparisons (Fig. 4A). Those observations match well with the in-gel fluorescence data described above where APP/PS1 tissue showed dramatically higher fluorescence labeling than the WT samples (Fig. 3C). Of all the prenylation substrate proteins enriched in the APP/PS1 mice, 15 of them were observed in all three comparisons. Those proteins are 4-trimethylaminobutyraldehyde dehydrogenase (Aldh9A1), Ras-related protein Ral-A (Rala), Synaptobrevin homolog Ykt6 (Ykt6), Ubiquitin carboxyl-terminal hydrolase isozyme L1 (Uchl1), Cell division control protein 42 homolog (Cdc42), Ras-related protein Rap-1b;1a,(Rab1b;Rap1a), 2,3-cyclic-nucleotide 3-phosphodiesterase (Cnp), Rab10, Rab11a; b, Rab18, Rab1a;b, Rab2a;b, Rab35, Rab3a, and Rab6a;b (Fig. 4C). Many of these proteins have previously been implicated as key players in the processing and regulation of Aβ peptide, APP, or tau proteins as highlighted in Table 1. It should also be noted that while there were a few proteins that were enriched in the WT mice compared with the APP/PS1 mice, none of those were observed consistently across three pairs.

**Figure 4.**
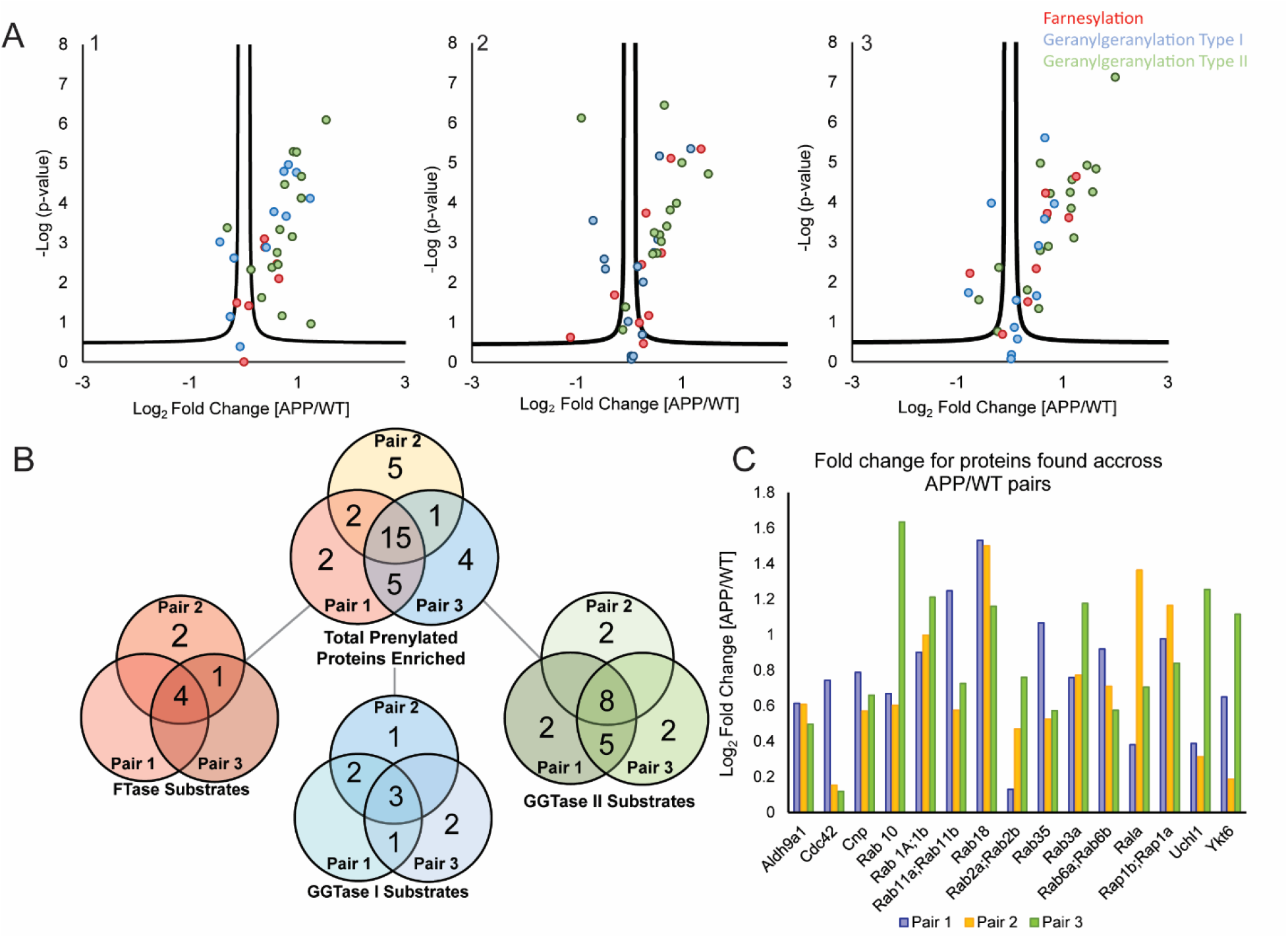
Prenylomic analysis of three pairs of APP/PS1 vs WT mice both subjected to ICV infusion of C15AlkOPP. (**A**) Volcano plots for each pair of mice (FDR=5% for all three plots). Data was processed in MaxQuant with the same parameters with non-prenlyated proteins removed for clarity. Color scheme: prenylated proteins that are known substrates for FTase (red); prenylated proteins that are substrates for GGTase I (blue); prenylated proteins that are known substrates for GGTase II (green). (**B**) Venn diagram showing the distribution of prenylated proteins across the three pairs of mice. This is further subdivided by the type of prenylation shown in color. Proteins that are grouped differently across the pairs are entered as one protein. Example: In pair 2 Hras;Kras;Nras are grouped, and in pair 3 Kras is not grouped and Hras;Nras are. So for simplicity, Hras;Nras and Kras are one data point in the Venn diagram. (**C**) Graph indicating the fold-change for the 15 proteins observed across all three pairs.

**Table 1:**
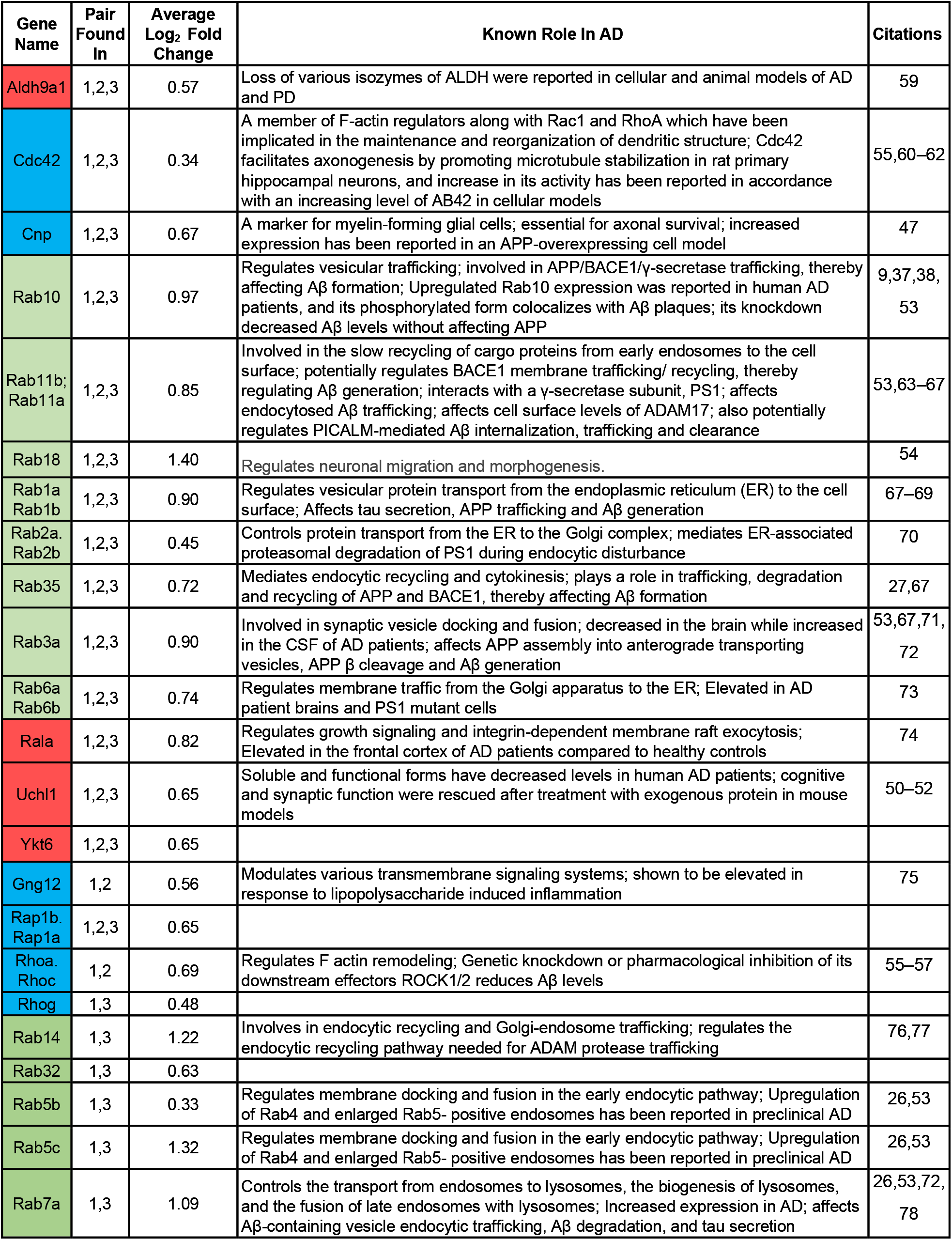
Proteins enriched in APP/PS1 Mice vs WT across all mice pairs. All proteins were found to be enriched in at least two pairs of mice. Red gene names are FTase substrates, green are GGTase II substrates, and blue are GGTase I substrates.

### Enrichment and protein-protein interaction analyses revealed overrepresented Gene Ontology (GO) terms and KEGG pathways among identified prenylated proteins and their interactions with AD network proteins

Finally, to gain additional insight into the functions of enriched prenylated proteins in the brains of APP/PS1 versus WT mice, functional enrichment analysis was performed using the complete list of all enriched proteins (36 total) including those that were found to be enriched only once, twice or in all three experiments (Fig. 5A). GO term enrichment analysis indicated endosomal transport (38%), cortical cytoskeleton organization (33%), and peptidyl-cysteine methylation (15%) as the top three biological processes, and synaptic vesicle membrane (47%) as the most enriched cellular component term (Fig. 5A). As expected, molecular function terms that are associated with GTPase activity were significantly enriched as well as Ras signaling (62%) in KEGG pathways (Fig. 5A). To investigate how these enriched prenylated proteins may be involved in AD pathogenesis, a protein-protein interaction network of these proteins and known AD network proteins^41,42^ was constructed (Fig. 5B). A total of 19 out of 36 prenylated proteins enriched in the brains of APP/PS1 compared to WT mice directly or indirectly interact with AD network proteins including amyloid precursor protein (APP) and microtubule-associated protein tau (MAPT). The roles of AD network proteins interacting with these prenylation substrates are summarized in Extended Data Table 1.

**Figure 5.**
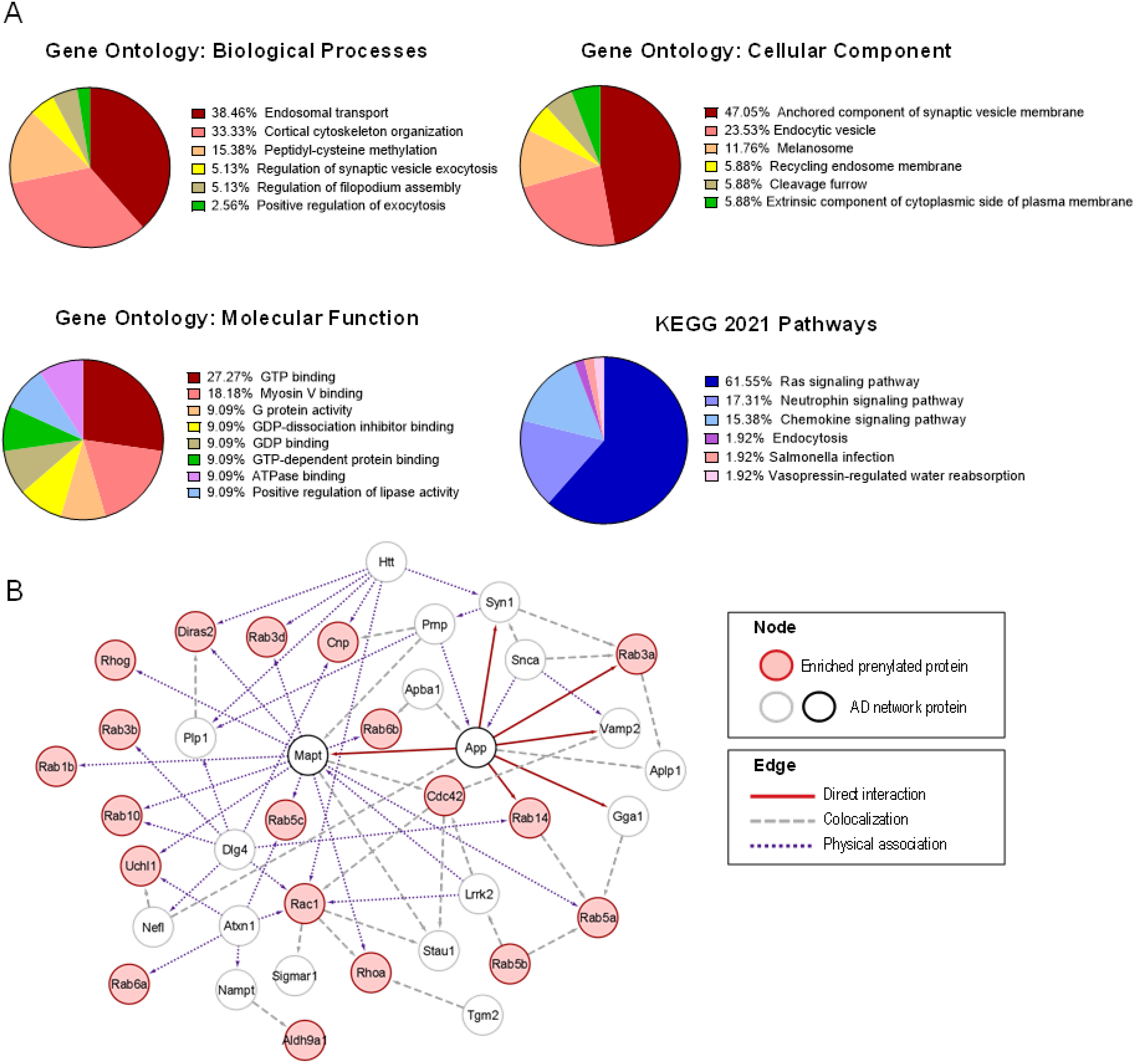
Gene Ontology (GO) term, KEGG pathway enrichment analyses and protein-protein interaction network of identified prenylated proteins and known AD network proteins. (**A**) Top enriched GO terms and KEGG pathways among prenylated proteins enriched in APP/PS1 mice. (**B**) Protein-protein interaction network showing direct/indirect interactions between AD network proteins and enriched prenylated proteins in APP/PS1 mice. A total of 19 prenylated proteins have known interactions with proteins involved in AD pathogenesis including amyloid precursor protein (App) and microtubule-associated protein tau (Mapt).

## Discussion

Despite the functional importance of protein prenylation in biological systems, experiments that monitor changes in the levels of multiple prenylated proteins, simultaneously, in whole animals have not been reported to date. In this study, the profiling of the brain prenylome in live mice was tested using a synthetic alkyne-modified isoprenoid analogue, C15AlkOPP. Recently, the same analogue was employed to characterize the prenylome in neuronal and glial cell lines as well as primary astrocytes,^35^ demonstrating its utility as a reporter for prenylation in multiple brain cell types. Due to the limited brain penetration of the analogue when administered via IP injection, a stereotaxic brain injection strategy was used to bypass the BBB. Although direct intracerebral injection would be preferable to introduce the analogue into the brain, this route is not suitable for a large injection volume or for global brain delivery. To minimize injection-associated damage while maximizing the delivery of the probe to different brain regions, the lateral ventricle was selected as the site of injection. Because of anatomical interconnectivity of the brain ventricular system, a single side ICV injection/infusion of C15AlkOPP was expected to result in metabolic labeling of both ipsilateral and contralateral sides of the brain. However, even with prolonged infusion (∼2 weeks), labeling was mostly restricted to the brain regions ipsilateral to the site of injection/infusion and further limited to the brain regions that are within ∼2 mm from the infusion site. *In situ* visualization of the labeling obtained with this probe indicated that a large fraction of the injected probe remained in the extracellular space in the brain parenchyma. Nevertheless, sufficient analogue incorporation was obtained using the ICV injection strategy to analyze the brain prenylome.

Consistent with previous findings,^35,38,39^ statin treatment augmented isoprenoid analogue incorporation. Metabolic labeling in the brain with C15AlkOPP resulted in the identification of 17 prenylated proteins in the absence of statin. Upon cotreatment with simvastatin, a total of 37 prenylated proteins were enriched including 8 farnesylated proteins, 9 GGTase I substrates and 20 GGTase II substrates. While the enhanced probe incorporation in the presence of the statin expanded the number of prenylated proteins that could be identified, we elected to perform subsequent labeling experiments with APP/PS1 AD model mice in the absence of statin reasoning that the presence of a statin could perturb cell physiology and complicate the interpretation of any differences observed between APP/PS1 and WT mice. Notably, *in vitro* labeling with C15AlkOPP was employed recently to probe the mouse brain prenylome in a related study. In contrast to the experiments performed here where analogue incorporation occurred *in vivo*, the *in vitro* labeling study used brain lysate obtained from mice engineered with neuron-specific knockouts of either FTase or GGTase.^43^ In this latter type of experiment, C15AlkOPP labeling was obtained by incubating brain lysate with C15AlkOPP and exogenous FTase or GGTase I followed by biotinylation, enrichment and proteomic analysis. Using that *in vitro* labeling approach, 13 farnesylated proteins and 7 GGTase I substrates were identified.^43^ It is interesting to note that only one of the proteins found enriched in this study were found enriched across all three APP/PS1 pairs (Rap1b;1a), and one (Rhoa), was found enriched in at least 2 pairs.^43^ Those differences probably reflect the different conditions those experiments were performed under. While the *in vitro* labeling strategy detects the unprenylated forms of proteins resulting from the genetic deletion of FTase or GGTase I in neurons the *in vivo* metabolic labeling detects prenylated proteins naturally occurring in the brain. Further, as the half-lives of prenylated proteins can be different from their unprenylated counterparts, these differences are not surprising and serve to highlight the utility of the *in vivo* approach described here.

Application of the ICV injection/infusion strategy detailed here comparing three pairs of APP/PS1 and WT mice yielded a total of 36 prenylated proteins enriched in the APP/PS1 mice across all three pairs. Of those 15 were seen in all three paired experiments with another 9 observed in at least two pairs. No prenylated proteins were consistently detected at higher levels in the WT mice compared with the APP/PS1 mice. Taken together, those results suggest that the levels of at least some prenylated proteins are elevated in AD. When comparing the APP/PS1 results to those of the initial prenylome profiling of WT mice it should be noted that four of the 15 commonly enriched proteins, Rab35, Rab3a, Cnp, and Ykt6,were found to have greater fold changes in the APP/PS1 mice than even in the statin treated mouse (Extended Data Fig. 4A) showing that the increase in prenylation of these proteins seen in the AD model surpasses the expected upper limit of enrichment established by the SV and C15AlkOPP profiling experiment. These results are consistent with the higher levels of C15AlkOPP incorporation observed in the in-gel fluorescence experiments reported above.

Importantly, many of the 15 prenylated proteins enriched in APP/PS1 mice have been previously implicated in human AD as summarized in Table 1. For example, mRNA expression of Rab10 was significantly increased in the temporal cortex of AD patients, and knockdown of Rab10 was shown to reduce Aβ42 levels.^9,44,45^ Also, the stress hormone induced downregulation of Rab35 has been shown to cause tau hyperphosphorylation. Overexpression of Rab35 in the hippocampus of mice was sufficient to prevent stress-induced tau accumulation and downstream dendrite and spine loss.^46^ Recently a direct role of Rab35 in the regulation of BACE1 activation and APP cleavage was reported.^27^ In addition, a SILAC-based proteomics study found elevated levels of a myelin protein Cnp in APP-overexpressing rat neuroblastoma B103 cells.^47^ Notably, the known prenylated protein Uchl1,^48,49^ which was found to be enriched in all experiments here, had not been reported in previous prenylomic experiments.^31,35^ Intriguingly, membrane-associated farnesylated Uchl1 promotes α-synuclein neurotoxicity and is implicated in the pathogenesis of Parkinson’s disease (PD).^48^ The down regulation and extensive oxidation of Uchl1 is observed in AD as well as PD patients; in particular in sporadic AD the levels of soluble cytosolic Uchl1 are inversely proportional to neurofibrillary tangle amounts.^50–52^ Furthermore, a subset of protein hits including Rab18 was consistently enriched in the APP/PS1 mouse.^53^ Since Rab18 is a known regulator of neuronal migration and structure formation,^54^ its augmented prenylation in APP/PS1 mice may perturb neuronal homeostasis and contribute to AD. Additional prenylated proteins enriched in APP/PS1 mice in two of the three replicate mouse pairs have also shown AD-associated changes in expression and signaling. Rhoa showed changes in expression level and its downstream signaling related to AD^55^ while genetic knockdown or pharmacological inhibition of the downstream effectors of Rhoa, ROCK1/2, have been shown to reduce Aβ levels as well as Aβ-induced cytoskeletal instability in neurons.^55–57^ Interestingly, when examining previously reported RNA-seq data comparing the mRNA levels in APP/PS1 mice to WT mice, no statistically relevant changes in the levels of the 15 proteins highlighted here was noted (Extended Data Fig. 3B).^12^ There were also no changes at the protein level for those same proteins when the APP/PS1 mouse proteome was profiled, (Extended Data Fig. 3A).^58^ It is interesting to note that the enzymes responsible for FPP and GGPP biosynthesis as well as the prenyltransferase enzymes themselves all showed no statistically significant dysregulation between the APP/PS1 mice and WT mice in either the aforementioned transcriptomic or proteomic studies (SI Table 2). Similar analysis of the enzymes in the mevalonate pathway, the biosynthetic source of isoprenoids showed no change in expression or abundance between the APP/PS1 mice and the WT except for one enzyme, isopentenyl-diphosphate delta-isomerase, which produces the key intermediate dimethylallyl diphosphate (DMAPP) that was found to be at much higher relative abundance in the proteomic analysis of APP/PS1 mice (SI Table 2). Such an increase could increase the pool of isoprenoid diphosphates available for protein prenylation. Indeed, higher levels of FPP and GGPP have been found in brain tissue samples obtained from AD patients.^20^

Bioinformatic analysis indicates that many of the enriched prenylated proteins are involved in cytoskeletal organization (Rho GTPases) and vesicle-mediated transport (Rab GTPases) which are important biological processes for synaptic plasticity and cellular component trafficking/recycling. Protein-protein interaction network analysis between these proteins and AD-associated proteins revealed that approximately half of the enriched prenylated proteins in the brain APP/PS1 mice directly or indirectly interact with AD pathology-related proteins. Those results suggest that dysregulated upregulation of prenylation of these proteins is potentially involved in pathogenic processing of APP and tau.

Overall, this study demonstrates the utility of metabolic labeling *in vivo* with an alkyne-modified isoprenoid analogue for studying disease-associated changes in prenylated proteins in AD. With prolonged infusion of the isoprenoid analogue into the cerebral ventricle, brain tissue was successfully labeled with the analogue in live mice. Through a subsequent proteomic analysis, fifteen prenylated proteins were found to be consistently enriched in APP/PS1 AD mice. These results demonstrate increased protein prenylation in AD, a new and potentially highly significant observation. The elevated levels of specific prenylated proteins noted here could potentially be useful for the development of new diagnostic tools for AD detection and the proteins identified pursued as novel therapeutic targets. Finally, the overall strategy reported here involving prolonged infusion, metabolic labeling, enrichment and quantitative proteomic analysis should be applicable to the study of other posttranslational modifications in mouse models of brain disorders.

## Methods

### Animals

Heterozygous APP/PS1 double transgenic mice on a mixed genetic background (B6xC3H) (B6C3-Tg (APPswe, PSEN1ΔE9) 85Dbo/J; stock# 004462) and their littermate wild type (WT) controls were kept under SPF conditions. Mice were PCR genotyped with DNA extracted from tail/ear biopsies and maintained on a 12 h-light/dark cycle with food and water ad libitum. All animal procedures used for this study were reviewed and approved by the Institutional Animal Care and Use Committee of the University of Minnesota (IACUC protocol # 1908-37310A).

### Preparation of analogue and simvastatin

Synthetic alkyne-modified analogue, C15AlkOPP was prepared as previously described.^30^ After dissolved in 25 mM NH_4_HCO_3_, the concentration was determined using ^31^P-NMR using a NaPO_4_ internal standard and the solution was sterilized by filtering through a 0.22-μm syringe filter followed by dilution to the desired concentration. Simvastatin (SV) (Calbiochem, 567020) was converted from the inactive lactone prodrug to the active acid form as previously described.^19^ In brief, 10 mg of SV powder was dissolved in 200 μL of ethanol (100%), and 163 μL of 1 N NaOH was added for alkaline hydrolysis. This stock solution was stored at −20 °C until use for up to 1 month. On the day of surgery, the stock solution was neutralized with 1 M HCl, diluted to 10 μM with sterile PBS, and filtered through a 0.22-μm syringe filter.

### Mouse surgery

Acute bolus intracerebroventricular (ICV) injection: For bolus injection, mice were deeply anesthetized with isoflurane throughout the surgery. 10 μL of vehicle (25 mM NH_4_HCO_3_), 10 μL of 150 mM C15AlkOPP, or 20 μL of 150 mM C15AlkOPP + 10 mM SV mixture (1:1) was loaded into a Hamilton syringe (Hamilton, 80300; 80501), and stereotaxically injected into the left ventricle at an administration rate of 1 μL per min using a microinjector using the following coordinates: A/P -0.5 mm, M/L +1.1 mm, D/V -2.5 mm. Brain tissues were collected 48 h after the injection. Chronic ICV infusion: The Brain infusion kit 3 (Cat# 0008851; Pump model #1002) was purchased from Alzet (Cupertino, CA, USA). For initial testing each pump was filled with one of the following: 100 μL of vehicle (25 mM NH4HCO3); 150 mM C15AlkOPP; 1:3 mixture of 10 mM simvastatin and 150 mM C15AlkOPP. For comparison between APP/PS1 mice and wild type controls, each pump was filled with 100 μL of 100 mM C15AlkOPP. A single 0.5 mm spacer was attached to the cannula to adjust the depth to 2.5 mm. A 2-cm long catheter tube was used to connect the osmotic pump with the cannula. The assembled brain infusion pump was placed in sterile normal saline until use. Pumps were filled and assembled in a class II biosafety cabinet to achieve sterility. Prior to and throughout the surgery, mice were deeply anesthetized by isoflurane. The osmotic pump was implanted into the subcutaneous space in front of the left hind limb, and the cannula was stereotaxically placed to the left ventricle using the same coordinate as the bolus injection. After 13 days, the pumps were removed, and the brains were harvested.

### Brain tissue collection and processing

Acute ICV injection and chronic ICV infusion comparison samples: After harvesting, the olfactory bulbs and cerebellum were removed from the brains. Then the brains were cut sagittally into the left and right hemispheres, both of which were acutely sliced coronally using a Vibratome (Leica Microsystems Inc.) at an alternating thickness of 400 μm and 600 μm from anterior to posterior. The 600 μm brain slices were fixed in 4% paraformaldehyde (PFA) at room temperature for 20 min, rinsed with PBS, and stored in 4°C immersed in PBS containing 0.02 % sodium azide until use. The 400 μm brain slices were homogenized in 250 μL of PBS containing 1% SDS and AEBSF using a Bullet Blender® tissue homogenizer (Next Advance, Inc., NY), then further homogenized by applying 2-s sonicating pulses 15 times with 4 s intervals on ice. Lysates were cleared by centrifugation at 13,000 x g for 15 min at room temperature, and protein concentrations of supernatants were measured by Bradford assay (Thermo Fisher, 23246) following the manufacturer’s protocol.

APP/PS1 and WT comparison samples: After removing the olfactory bulbs and the cerebellum, the brains were cut sagittally into left and right hemispheres. About 3 mm-thick brain slices (coronal) around the infusion site were dissected out, homogenized in 800 μL of PBS containing 1% SDS and 1 mM AEBSF using a Bullet blender, and sonicated with 2-s pulses 15 times with 4 s intervals on ice. Lysates were cleared by centrifugation at 13,000 x g for 15 min at room temperature, and the protein concentrations were determined by Bradford assay.

### Click reaction and in-gel fluorescence

Lysates were diluted to 1μg/μL protein for click reaction. Click reaction was carried out by sequentially adding TAMRA-PEG4-N3 (25 μM, BroadPharm), tris(2carboxyethyl)phosphine (TCEP, 1 mM, Sigma-Aldrich), tris(1-benzyl-1H-1,2,3-triazol-4-yl-)methylamine (TBTA, 100 μM, Sigma-Aldrich), and CuSO_4_ (1 mM, Sigma-Aldrich) with vortexing, and incubating at room temperature for 90 min on a multi-tube rotator. After the reaction, proteins were precipitated using a ProteoExtract® protein extraction kit (MilliporeSigma) following the manufacturer’s protocol. Protein pellets were washed, air-dried, and dissolved in Laemmli buffer (2% SDS, 10% glycerol, 62.5 M Tris HCl, pH 6.8, 0.01% bromophenol blue, and 5 % β-mercaptoethanol). After heating at 100 °C for 5 min, samples were resolved using a 12% SDS-PAGE gel. TAMRA fluorescence in gels was detected using an iBrightTM imaging system (Invitrogen).

### Immunoblot analysis

Brain tissue lysates were subjected to 12% SDS-PAGE under the reducing condition, and proteins were transferred to a PVDF membrane. The membrane was incubated with 1:1000 rabbit anti-β-APP antibody (Invitrogen, 51-2700) overnight at 4°C, then with 1:3000 HRP-conjugated mouse anti-rabbit antibody (Santacruz Biotech, sc-2357). After antibody incubations, membranes were treated with the Clarity Western ECL substrate (Bio-Rad, 1705061). Signals were captured using an iBrightTM imaging system (Invitrogen).

### In situ click reaction of brain sections and fluorescence imaging

Fixed 600 μm-thick brain slices were further sectioned to a thickness of 50 μm after embedded in 3% agarose. Sections were permeabilized and blocked by incubating in the permeabilization buffer (0.5 % Triton x-100, 5 % normal donkey serum, 1 % bovine serum albumin (BSA) in PBS (pH 7.8)) at room temperature overnight, then washed with 0.1 % Triton X-100 in PBS 3 times. The click reaction mixture was assembled by sequentially adding 2 μM TAMRA-PEG4-N3, 0.5 mM TCEP, 0.1 mM TBTA, and 0.2 mM CuSO_4_. Brain sections were incubated in the click reaction mixture in the dark with gentle shaking at room temperature overnight. Brain sections were incubated in the click reaction mixture in the dark with gentle shaking at room temperature overnight. Any remaining click reaction mixture on the brain sections was extensively washed with the click reaction wash buffer (1% Tween-20, 0.5 mM EDTA-containing PBS (pH 7.8)) for 20 min, 3 times, then with PBST (0.05% Tween-20 in PBS (pH 7.8)) for 10 min, 2 times. Sections were then incubated in 1:400 rabbit anti-Neu-N antibody (Cell Signaling Technology, 24307). The next day, sections were washed 3 times with PBST, 2 times with PBS, and incubated with 1:500 Alexa 488-conjugated donkey anti-rabbit antibody (Invitrogen, A21206) for 2 h at room temperature. After washing 3 times with PBST and 2 times with PBS, sections were mounted on microscopic slides with ProlongGold antifade mountant with DAPI (Thermo Fisher, P36935), and imaged with a Keyence All-in-one fluorescence microscope (BZ-X800E).

### Enrichment of labeled proteins and sample preparation for proteomics

Protein concentrations of brain tissue lysates were determined with a BCA assay (Bio-Rad). Then 2 mg in lysis buffer from each treatment was subjected to click reaction with biotin-N3 (BroadPharm) by sequentially adding 100 μM, biotin-N3 (10 µL), 1 mM TCEP (20 µL), 0.1 mM TBTA (10 µL), and 1 mM CuSO_4_ (20 µL). All samples were then diluted to 1 mL with PBS containing 1% SDS, followed by a 120-min incubation at room temperature in the dark. Proteins were precipitated out using chloroform–methanol (1:4:3) extraction, and protein pellets were washed with methanol. Pellets were air dried and were then stored at -20 ^°^C overnight followed by dissolution in 0.55 mL PBS containing 1% SDS. Protein concentrations were measured using a BCA assay. Biotinylated samples (1.2 mg of protein) were incubated with pre-washed Neutravidin agarose beads (200 µL, 50% slurry, Thermo Scientific) in 500 µL of PBS containing 1% SDS for 120 min, then beads were washed 3x with 1 mL PBS + 1% SDS, PBS, 3x 1 mL 8 M urea, and 3x 1 mL 50 mM triethylammonium bicarbonate (TEAB). Resin was resuspended in 100 µL of 50 mM TEAB buffer, and proteins were digested on-bead with trypsin (1.5 µg, in provided buffer, Promega Corp.) overnight at 37 °C, and peptides were collected from beads using spin columns by washing with 50 mM TEAB and dried by lyophilization. Peptides were dissolved in 50-60 µL and their resulting concentrations determined via BCA assay. Three 10 μg aliquots of peptides from each mouse were used as technical replicates. Samples were supplemented with 150 fmol of internal standard (yeast ADH1, Waters) and subjected to tandem mass tag (TMT)-labeling using TMT 6-plex reagents (Thermo Fisher Scientific). TMT 126-128 were used for the reference (Vehicle or WT) and TMT 129-131 were used for the APP/PS1 or C15AlkOPP WT samples. TMT-labeled samples were combined, dissolved in 200 mM NH_4_HCO_2_ (pH 10, 300 µL), then subsequently fractionated using a homemade stage tip (three layers of SDB-XC extraction disks, (3M, 1.07 mm x 0.50 mm i.d.)) in a 200 µL pipette tip. Peptides were then fractionated under high pH reverse phase conditions yielding 7 fractions of 60 µL in volume (5, 10, 15, 20, 22.5, 27.5, 80% CH3CN in 200 mM NH_4_HCO_3_, pH 10). The first two fractions were combined, then all fractions were dried by lyophilization. Samples were then dissolved in 30 µL of 0.1% formic acid for LC-MS^3^ analysis. This method is based on a previously reported method.^35^

### LC-MS data acquisition

Data acquisition was performed as described previously.^35^ Briefly, TMT-labeled peptides were resolved at a flow rate of 300 nL/min using a RSLC Ultimate 3000 nano-UHPLC (Dionex) in a reversed-phase column (75 μm i.d., 45 cm) packed in-house.^79^ Each fraction from the high pH prefractionation was subjected to varying gradients (7-34 %) of acetonitrile (with 0.1 % formic acid) in solvent (0.1% formic acid in water) for 80 min and sprayed directly into the Orbitrap instrument (Thermo Fisher Scientific). MS1 scans were collected at 120,000 resolutions in a 320–2,000 m/z range with 100 ms max injection time (IT) and automatic gain control (AGC) target of 200,000. The subsequent data-dependent MS/MS scans were collected with collision-induced dissociation (CID) at a normalized collision energy (NCE) of 35% with a 1.3 m/z isolation window, with max IT of 100 ms and AGC target of 5000. Acquisition at MS3 was done by selecting the top 10 precursors (SPS) for fragmentation by high-collisional energy dissociation (HCD) in the orbitrap with the following settings: 55% NCE, 2.5 m/z isolation window, 120 ms max IT, and 50,000 AGC target.

### Prenylomic Data processing

Raw data files were uploaded into MaxQuant (version 1.6.17.0) and searched against a non-redundant mouse database (UP0000000589) from Uniprot. For all but the following parameters, the default settings were maintained. Trypsin/P was used for digestion with allowance for 3 missed cleavages, minimum peptide length was set to 7, protein FDR was set to 0.5, modifications in search oxidation (M) and Acetyl (protein N-Term), and unique and razor peptides were used for quantification. MaxQuant was run through the MaxQuantCmd.exc at the University of Minnesota Supercomputing Institute.

The proteingroup.txt file generated was uploaded into Perseus (version 1.6.14.0). Proteins that were potential contaminants and reverse peptides were removed by site identification. Raw intensities were transformed to log_2_ values, and proteins with more than 3 out of 6 values returning “NaN” after transformation were removed. Missing values were imputed from the normal distribution of the remaining values. Reported values (TMT) were normalized by subtracting rows by mean value and columns by median value. Statistical analysis was performed using a two-sample t-test FDR = 5 % and s0 = 0.1. Data was exported to excel for generation of plots and figures.

### Gene Ontology and KEGG pathway enrichment analyses and interaction network construction for identified prenylated proteins and AD network proteins

Functional enrichment and protein-protein interaction analyses were performed using the complete list of 36 prenylated proteins identified in any of the three pair comparisons. Grouped isoforms that cannot be distinguished from each other with mass spectrometry due to high peptide sequence homology were separated into different isoforms to facilitate the analyses. Gene Ontology and KEGG pathway enrichment analyses for the enriched prenylated proteins in APP/PS1 mice were performed in Cytoscape software (version 3.9.0) using the ClueGO V2.5.8 plug-in. Two-sided hypergeometric tests were used to calculate the significance of GO terms or KEGG pathways, and a P-value less than 0.05 with Benjamini-Hotchberg adjustment were considered statistically significant. The protein-protein interaction (PPI) network was constructed using the Biological General Repository for Interaction Datasets (BioGRID) in Cytoscape software (version 3.2.1).^42^ Then prenylated proteins enriched in APP/PS1 mice were mapped along with proteins in the previously curated AD network by Brueza *et al*.^41^ on the mouse PPI network. A subnetwork was constructed by selecting AD network proteins with known direct or indirect interactions with the enriched prenylated proteins.

## Extended Data Figures

**Extended Data Figure 1.**
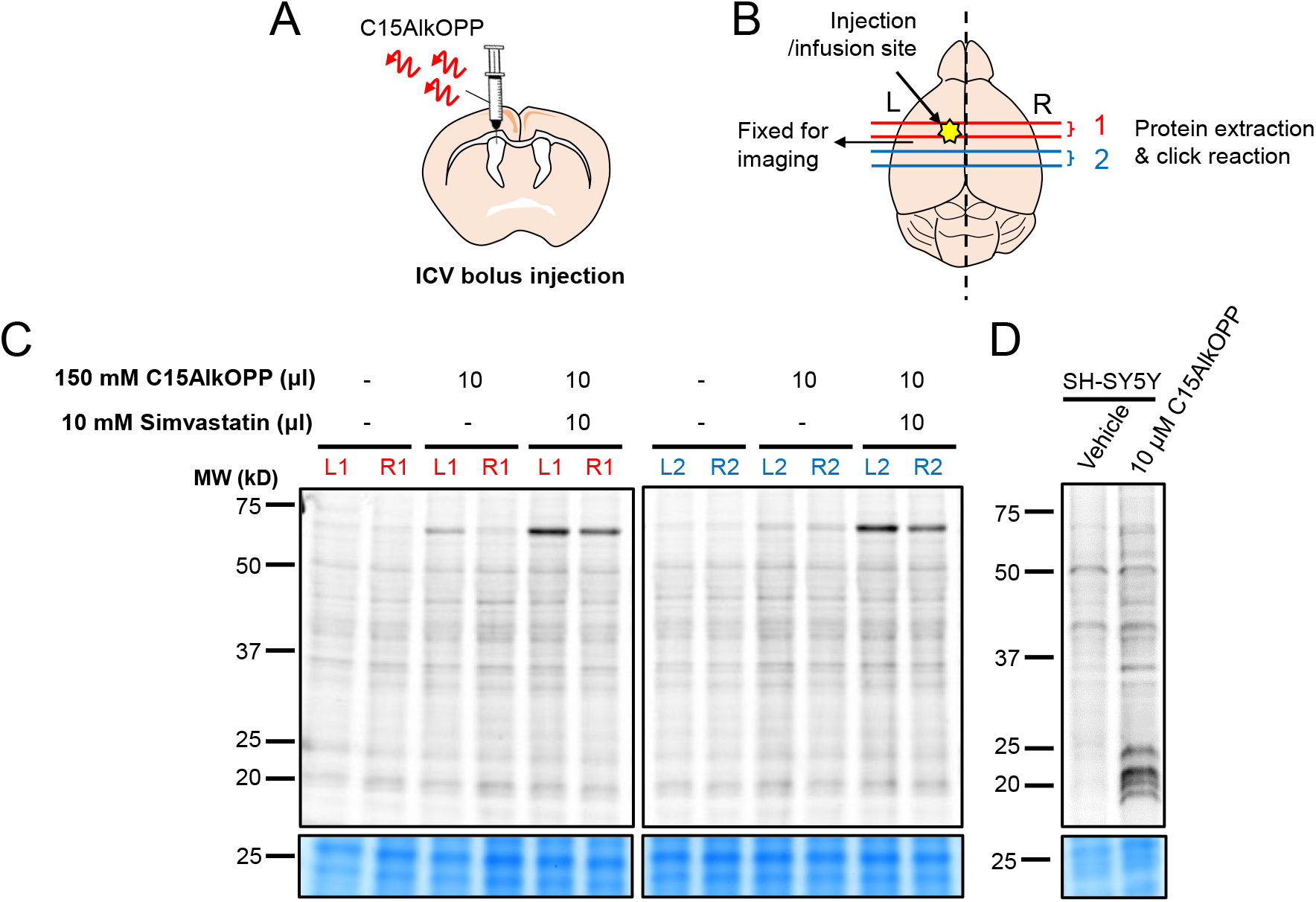
A single bolus ICV injection of C15AlkOPP showed limited brain metabolic labeling *in vivo*. (**A**) Schematic representation of ICV bolus injection and (**B**) the locations of two brain coronal sections (400 μm) relative to the injection site that were used for the representative in-gel fluorescence. (**C**) In-gel fluorescence and Coomassie blue-stained gel images of brain tissue samples obtained from the regions depicted in (**B**). The brain section (600 μm) located between section #1 and #2 was PFA-fixed for *in situ* click reaction and imaging. Brains were harvested 48 hours after an ICV bolus injection of vehicle (25 mM NH_4_HCO_3_, 10 μL), 150 mM C15AlkOPP (10 μL), or 150 mM C15AlkOPP (10 μL) + 10 mM SV (10 μL). (**D**) In-gel fluorescence and Coomassie blue gel staining of human neuroblastoma SH-SY5Y cell lysate after 24-hour incubation with vehicle or 10 μM C15AlkOPP, diluted from the same analogue used for the *in vivo* study.

**Extended Data Figure 2.**
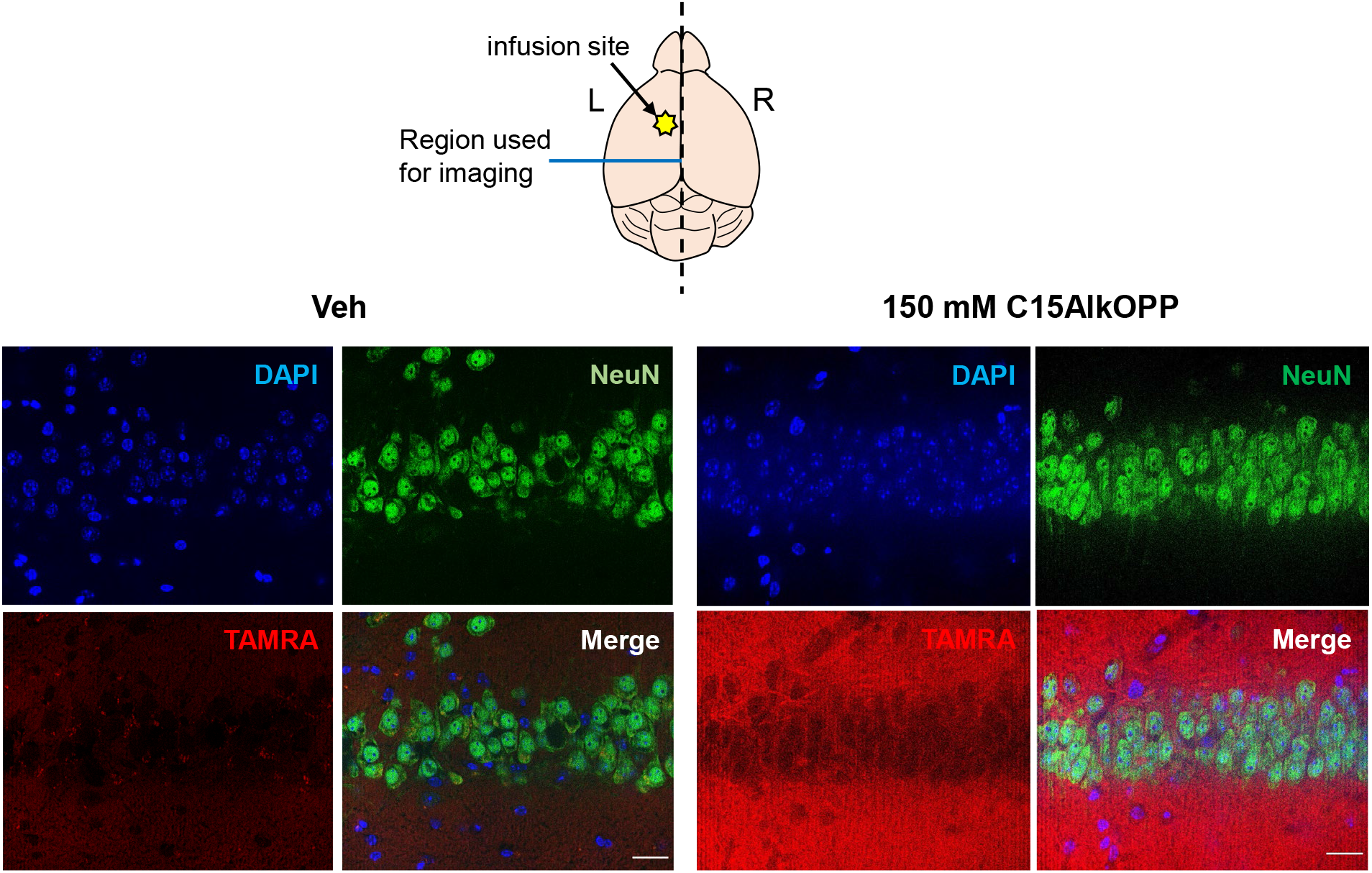
Visualization of C15AlkOPP incorporation in a brain section after in situ click reaction with TAMRA-azide. After 13-day left ICV infusion of 150 mM C15AlkOPP, tissue sections (50 μm) were fixed and subjected to *in situ* click reaction with TAMRA-azide along with NeuN immunostaining staining and DAPI staining. Top: Schematic representation of brain showing site of ICV infusion and region analyzed by in situ click reaction with TAMRA-azide. Left: Fluorescence microscopy of a brain section showing neurons (green), nuclei (blue) and TAMRA staining after ICV infusion with vehicle alone. Right: Fluorescence microscopy of a brain section showing neurons (green), nuclei (blue) and TAMRA staining after ICV infusion with 150 mM C15AlkOPP. Images were recorded at 40 × magnification on Keyence All-in-one fluorescence microscope (BZ-X800E). Scale bar 40 μm.

**Extended Data Figure 3:**
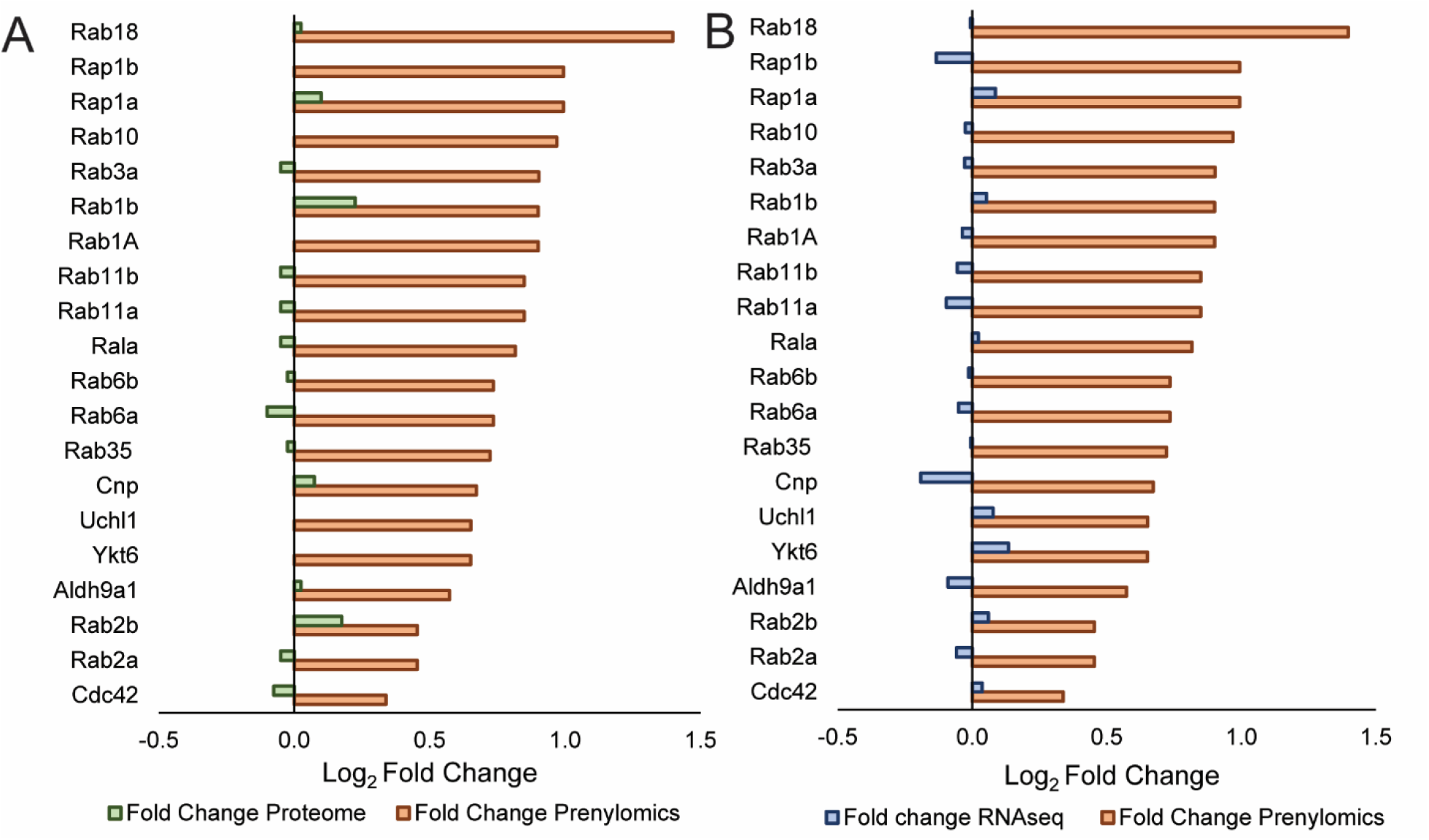
Transcriptomic and native abundance comparisons showed that prenylation is upregulated in the APP/PS1 mice. **A**) comparisons of the log2 fold change for the 15 common enriched proteins found in the prenylomic analysis of APP/PS1 mice (orange) compared to a total bottom-up proteomic analysis of APP/PS1 mice compared to WT from externally reported data from proteome exchange (green). **B**) comparisons of the log_2_ fold change for the 15 common enriched proteins found in the prenylomic analysis of APP/PS1 mice (orange) compared to previously reported RNAseq data comparing the expression of proteins in APP/PS1 mice to WT mice.

**Extended Data Figure 4:**
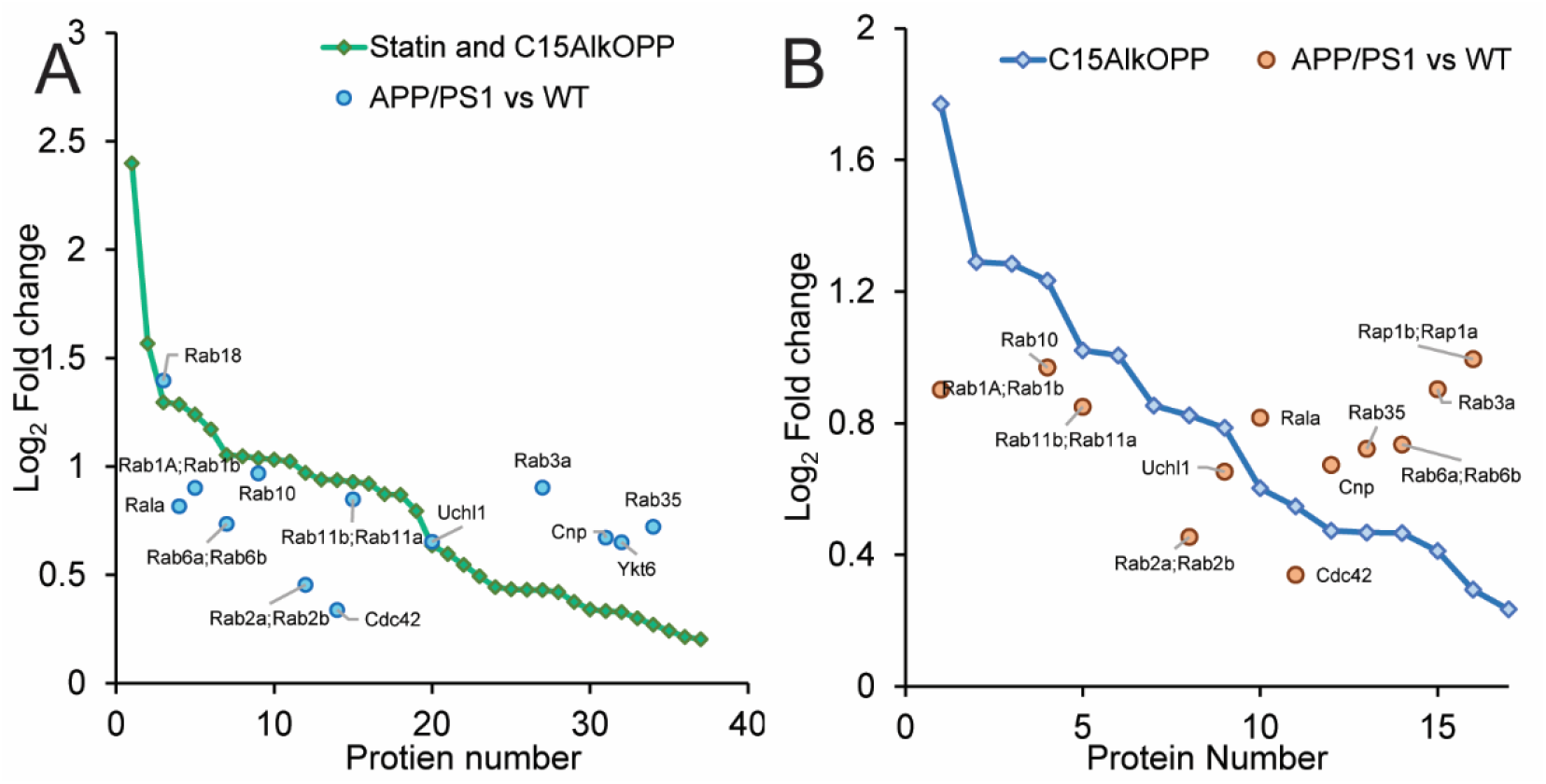
Ranking prenylated proteins found in profiling experiments by fold change and comparisons to the Fold change of the 15 common proteins. In both graphs, the proteins found enriched in either the C15AlKOPP and statin profiling experiment (**A**) or the C15AlkOPP profiling experiment (**B**) were ranked by decreasing Log2 fold change. **A**) All 15 in common proteins were found in the statin and C15AlkOPP profile, and their fold changes were overlaid with the ranking from the profiling experiment. Most proteins were found below the ranking line, as expected since the APP/PS1 mice did not receive the labeling enhancing statin treatment. However, some proteins were found above the ranking line suggesting that they are dramatically over prenylated in the APP/PS1 mice, (Rab3a, Cnp, Ykt6, Rab35). **B**) A similar analysis as (**A**) but with the 12 proteins found in the APP/PS1 pairs of mice that were also found in the C15AlkOPP profile. Rab 35 was found again to be prenylated more in the APP/PS1 mice along with the additional proteins Rab 6a, 6b and Rala, Cnp, Rab3a, Rap1b and Rap1a.

**Extended Data Table 1:**
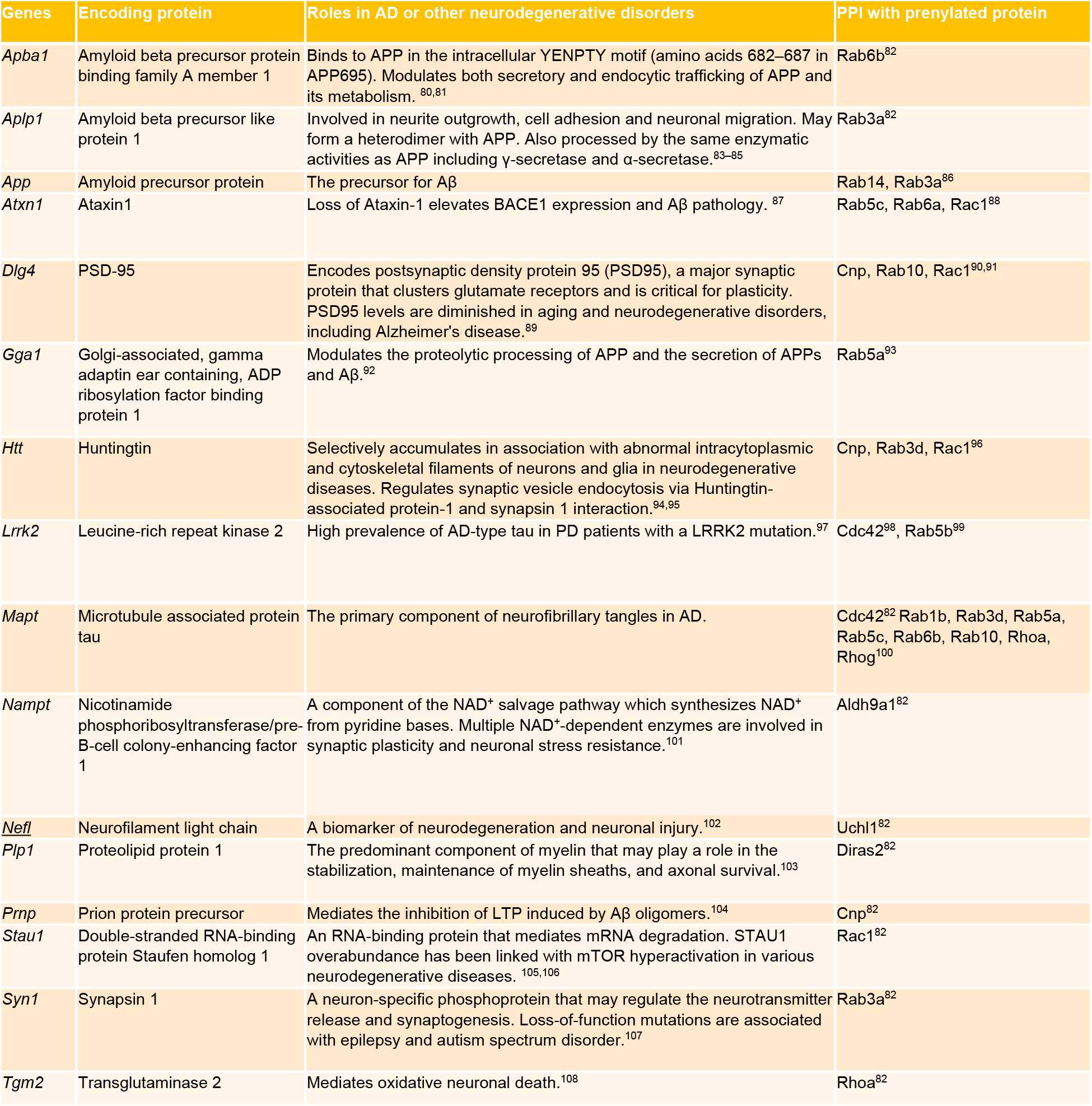
Protein-protein interaction analysis of prenylated proteins in dataset and AD network proteins.

## Data Availability

The mass spectrometry proteomics data have been deposited to the ProteomeXchange Consortium via the PRIDE partner repository with the dataset identifier PXD032350. The previously reported APP/PS1 RNAseq Data is deposited in Gene Expression Omnibus (accession# GSE 174314). The previously reported total proteome profiling of APP/PS1 mice was pulled from the ProteomeXchange Consortium via the PRIDE repository (accession# PXD015335).

## Acknowledgements

The authors acknowledge the Minnesota Supercomputing Institute (MSI) at the University of Minnesota for providing resources that contributed to the prenylomic research results reported within this paper (http://www.msi.umn.edu) and Dr. Yingchun Zhao and Dr. Peter Villalta for the assistance with proteomic data collection in the Analytical Biochemistry Shared Resource of the Masonic Cancer Center, designated by the National Cancer Institute and supported by P30 CA077598

## Author information

### Contributions

SA performed the prenylomic sample preparation and prenylomic analysis, and data processing.

AJ performed animal surgery, brain tissue collection, in-gel fluorescence, *in situ* imaging, functional enrichment and protein-protein interaction analyses.

LL and MD conceptualized, advised, and guided project development.

SA and AJ wrote the paper with input and guidance from all authors.

SA, AJ, LL, MD all interpreted data.

SM synthesized the C15AlkOPP probe for this study.

All authors critically read and reviewed manuscript.

### Additional Information

**Correspondence and requests for materials** should be addressed to Mark D. Distefano or Ling Li.

