## Supplemental Figures for "*In vivo* prenylomic profiling in the brain of a transgenic mouse model of Alzheimer’s disease reveals increased prenylation of a key set of proteins"

List of Figures and Tables:

1. SI Figure 1: Volcano plots of APP/PS1 vs WT mouse pairs showing all enriched proteins
2. SI Table 1: Values for RNAseq fold-change and native abundance fold-change compared to prenylomic fold-change of common ungrouped proteins found in APP/PS1 vs WT analysis
3. SI Table 2: $Log_2$ fold-change of enzymes related to prenylation from either proteomic profiling of APP/PS1 vs WT mice, or transcriptomic analysis of APP/PS1 vs WT mice

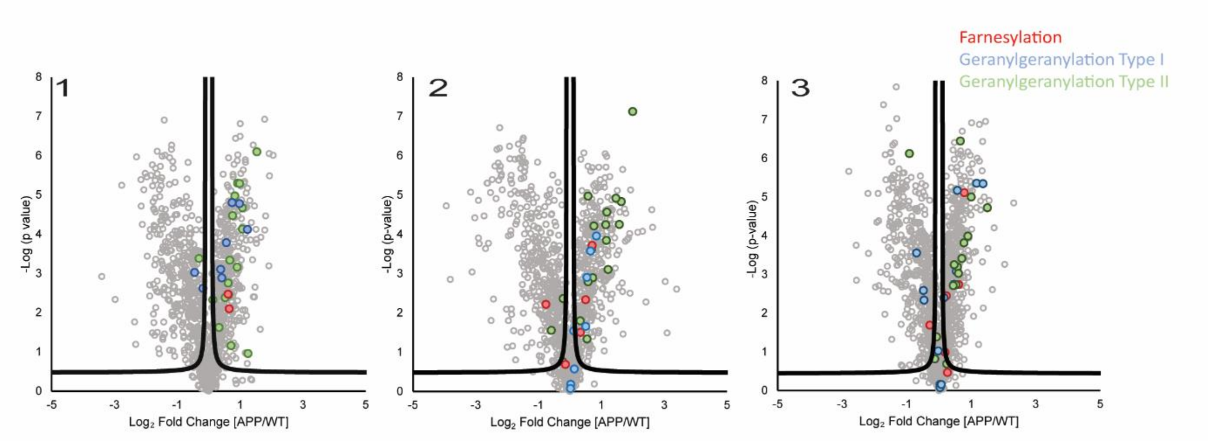


**SI Figure 1:** Volcano plots of data from APP/PS1 vs WT mouse pairs showing all enriched proteins. This figure presents the same data shown in Figure 4 in the main text with the exception that all enriched proteins including those not known to be prenylated are also included. Color scheme: prenylated proteins that are known substrates for FTase (red); prenylated proteins that are substrates for GGTase I (blue); prenylated proteins that are known substrates for GGTase II (green); proteins that are not known to be prenylated (grey).

| Gene Name | Fold Change $Log_2$ Prenylomics | Fold Change $Log_2$ RNAseq | Fold Change $Log_2$ Proteome |
|---|---|---|---|
| Aldh9a1 | 0.573 | -0.09 | 0.025 |
| Cdc42 | 0.339 | 0.037 | -0.075 |
| Rab 11a | 0.849 | -0.096 | -0.050 |
| Rab10 | 0.969 | -0.025 | 0.000 |
| Rab11b | 0.849 | -0.056 | -0.050 |
| Rab18 | 1.397 | -0.007 | 0.025 |
| Rab1a | 0.901 | -0.038 | 0.000 |
| Rab1b | 0.901 | 0.054 | 0.225 |
| Rab2a | 0.454 | -0.059 | -0.050 |
| Rab2b | 0.454 | 0.06 | 0.175 |
| Rab35 | 0.722 | -0.005 | -0.025 |
| Rab3a | 0.903 | -0.029 | -0.050 |
| Rab6a | 0.735 | -0.051 | -0.100 |
| Rab6b | 0.735 | -0.013 | -0.025 |
| Rala | 0.817 | 0.023 | -0.050 |
| Uchl1 | 0.652 | 0.077 | 0.000 |
| Ykt6 | 0.651 | 0.000 | 0.000 |
| Cnp | 0.673 | -0.191 | 0.075 |
| Rap1b | 0.994 | 0.086 | 0.000 |
| Rap1a | 0.994 | -0.133 | -0.025 |

**SI Table 1:** Values for RNAseq $Log_2$ fold-change[1] and proteomic $Log_2$ fold-change[2] compared to prenylomic fold-change for proteins found in APP/PS1 vs WT analysis, proteins are ungrouped for simplicity. Color scheme: prenylated proteins that are known substrates for FTase (red); prenylated proteins that are substrates for GGTase I (blue); prenylated proteins that are known substrates for GGTase II (green).

| Protein Name | Gene Name | $Log_2$ Fold-Change (RNAseq) | $Log_2$ Fold-Change (Proteomics) |
|---|---|---|---|
| Mevalonate kinase | Mvk | 0.21 | 0 |
| Diphosphomevalonate decarboxylase | Mvd | 0.21 | 0 |
| Phosphomevalonate kinase | Pmvk | 0.090 | -0.25 |
| Isopentenyl-diphosphate Delta-isomerase 1 | Idi1* | -0.099 | 33 |
| Farnesyl pyrophosphate synthase | Fdps | 0.15 | -0.05 |
| Geranylgeranyl pyrophosphate synthase | Ggpps | NA | -0.05 |
| Rab proteins geranylgeranyltransferase component A 1 | Chm (REP1) | -0.14 | 0 |
| Rab proteins geranylgeranyltransferase component A 2 | Chml (REP2) | -0.32 | NA |
| Prenylcysteine oxidase-like | Pcyox1l | 0.024 | 0.025 |
| Prenylcysteine oxidase | Pcyox1 | 0.057 | 0.13 |
| CAAX prenyl protease 1 homolog | Zmpste24 | -0.055 | -0.025 |
| Protein-S-isoprenylcysteine O-methyltransferase | ICMT* | -0.59 | NA |
| CAAX prenyl protease 2 | RCE1 | 0.062 | NA |
| Geranylgeranyl transferase type-2 subunit alpha | Rabggta | 0.0816 | -0.075 |
| Geranylgeranyl transferase type-2 subunit beta | Rabggtb | -0.068 | 0 |
| Protein farnesyltransferase subunit beta | Fntb | -0.34 | 0.25 |
| Protein farnesyltransferase/geranylgeranyltransferase type-1 subunit alpha | Fnta | -0.12 | -0.05 |

**SI Table 2:** $Log_2$ fold-change of enzymes related to prenylation from either proteomic profiling of APP/PS1 vs WT mice, or transcriptomic analysis of APP/PS1 vs WT mice. In red are proteins related to the mevalonate pathway, green are Rab escort proteins, grey are prenyl recycling enzymes, blue are post-prenylation processing proteins, brown are prenyltransferase enzyme subunits. *Only two enzymes were found to be statistically different: Idi1 was found to be upregulated in the APP/PS1 proteome, and ICMT was shown to have decreased mRNA expression in the APP/PS1 mice.
